# The ecological and economic benefits of sustainable agricultural practices: Evidence from on-farm trials in broad-acre grain crops

**DOI:** 10.1101/2024.10.21.619338

**Authors:** Luis Mata, Leo McGrane, James Maino, Grant Sims, Craig Drum, Paul A. Umina

**Author notes:** Corresponding authors: Luis Mata and Paul A. Umina.

## Abstract

1. Sustainable agricultural practices are essential for mitigating the negative environmental impacts of conventional agriculture while ensuring food security. However, widespread adoption of these practices requires robust evidence of their efficacy and economic viability.
2. We co-designed a two-year field trial with farmers and agronomy advisors in Australia, to evaluate the ecological and economic benefits of sustainable agricultural practices for managing the redlegged earth mite, a major pest of Australian grain crops. We compared ‘Novel’ treatments representing long-term farmer-implemented sustainable agricultural practices based on biological control with ‘Conventional’ treatments and ‘Plus’ treatments designed as counterfactuals to disentangle the effects of specific pest control and plant nutrient components.
3. Redlegged earth mite densities remained below economic thresholds across all treatments and years, demonstrating effective pest control in both conventional and sustainable systems. Notably, the Novel treatment supported higher densities of beneficial arthropods, indicating increased biological control potential.
4. Yield and gross profit margins were generally similar between the treatments, indicating that sustainable agriculture practices can maintain profitability while fostering biodiversity. Practical implication. Our study provides evidence that biological control and biofertiliser supplementation can be effectively used to manage agricultural pests and demonstrates the value of close collaboration with farmers and agronomy advisors to conduct ecological field research that has practical applications.

## Introduction

Agricultural intensification is essential to meet growing demands for food and fibre (Godfray et al. 2010; Giller et al. 2021) and, fundamentally more importantly, to rise to the ‘End hunger’ challenge set forth by the United Nations’ Sustainable Development Goals (United Nations 2015). Unfortunately, however, the overreliance on chemicals in intensive agroecosystems, particularly those used to control pests (Food and Agricultural Organization 2020), has negatively affected natural and anthropogenic systems, leading to residue, bioaccumulation, runoff and spillover processes affecting people (Rohr et al. 2019; Lefèvre-Arbogast et al. 2024), as well and other species (Hallmann et al. 2014; Jacobsen and Hjelmsø 2014; Zioga et al. 2023). On the farm level, these adverse impacts translate specifically into (1) acute toxicity, sub-lethal and transgenerational effects on non-target arthropods, including beneficial species providing biological control (Desneux et al. 2007; Schulz et al. 2021; Schmidt-Jeffris 2023; Mata et al. 2024); (2) emergence of secondary pest outbreaks (Hill et al. 2017); and (3) evolution in pests of pesticide resistance (Hawkins et al. 2019; Umina et al. 2019). This evidences a challenging conundrum aptly termed ‘The Pesticide Paradox’ (Enserink et al. 2013), in which pesticides have enabled the production of food and clothing and protection against zoonotic diseases while simultaneously posing profound threats to humanity and the rest of nature.

Ecological and sustainability intensification (Garnett et al. 2013; Kleijn et al. 2019) – as represented by an array of agricultural systems such as agroecology (Ewert et al. 2023), integrated pest management (Deguine et al. 2021), organic farming (Seufert et al. 2017), pesticide-free crop production (Finger and Möhring 2024), regenerative agriculture (O’Donoghue et al. 2022) and precision agriculture (Gebbers and Adamchuk 2010) – champion a more judicious and strategic approach to arthropod pest management. By adopting on-farm practices aimed at reducing or eliminating agrochemical applications, these systems can greatly contribute to ameliorating, stopping and reversing the problems caused by conventional farming (Bengtsson et al. 2005; Kremen et al. 2012; Choudhary et al. 2018; Muneret et al. 2018). Importantly, research has shown that ecological and sustainability intensification can be achieved while maintaining or increasing yields (Macfadyen et al. 2014; Pywell et al. 2015; Pecenka et al. 2021; MacLaren et al. 2022) and ensuring profitability for farmers (Lechenet et al. 2017; Jat et al. 2021; Gong et al. 2022).

Not surprisingly, sustainable agriculture practices are increasingly being used by farmers and recommended by agronomy advisors interested in diversifying their traditional regimes (Muhie 2022; Rehman et al. 2022). An essential component of the sustainable agriculture practice toolkit is biological control – the intrinsic pest and disease protection provided by a diverse range of beneficial organisms from viruses, bacteria and fungi to predatory and parasitoid arthropods (Bale et al. 2008). Farmers put biological control into gear by either conserving and promoting existing beneficials in non- and inter-crop habitats (Thomson and Hoffmann 2009; Alarcón-Segura et al. 2022) or augmenting their populations through the introduction of commercially reared individuals (van Lenteren et al. 2018). Important synergies can be achieved when biological control is aligned with other sustainable practices; for example: (1) applying integrated nutrient management principles such as the use of biofertilisers to complement the plant nutritional needs provided by synthetic fertilisers (Wu and Ma 2015; Kumar et al. 2022); (2) complementing chemical weed control with mechanical, thermal and cultural methods (Hatcher and Melander 2003); and (3) using remote sensing and information technologies to minimise input utilisation (Gebbers and Adamchuk 2010).

Farmers’ initial adoption and ongoing uptake of biological control and other synergistic sustainable practices, however, has been challenging due to a complex interplay of multi-faceted values, beliefs, attitudes and motivations. Uptake of sustainable innovations comes more naturally, for example, to farmers with ecocentric or pluricentric worldviews (Pascual et al. 2023), living by a non-utilitarian moral duty to preserve biodiversity and the environment (Klebl et al. 2024). Accordingly, farmers who believe that reducing their pesticide use will benefit nature are more likely to decrease their chemical usage (Bakker et al. 2021). Other intrinsic factors influencing adoption and uptake in farmers include their self-awareness of the genotoxic, chronic diseases associated with occupational exposure to pesticides (Silva et al. 2020); their concern that society would hold them responsible for biodiversity loss (Busse et al. 2021); and their inherit sense of responsibility to steward the land’s heritage (Sorice et al. 2023). Beyond these, a set of market-driven factors determine how farmers engage with sustainable practices. For example, there is increasing consumer demand for pesticide-free products (Nitzko 2024) and a growing need to comply with regional, national and international mandates, including agrochemical bans (Schebesta and Candel 2020; Gabriela et al. 2022), pesticide residue legislation (Nader et al. 2020), and insecticide resistance management strategies (Umina et al. 2019; Shaw et al. 2023). Of relevance are the recent policies enacted by the European Union, which aim to reduce by 50% the use of and risk from pesticides and to increase the area farmed organically to 25% by 2030 (European Union 2020a, 2020b).

Ultimately, farmers’ full and sustained uptake of sustainable pest management practices will be determined by a detailed understanding of whether they are having positive effects on crop production and, critically, are doing so while remaining economically viable. This knowledge can be obtained by researchers – collaborating closely with farmers and their advisors – conducting well-designed ecological experiments (Ockendon et al. 2021) and economic analyses (Naranjo et al. 2015). While interest in providing scientific evidence that sustainable agricultural practices are meeting the desired outcomes is high (Gurr and Wratten 2000; Kleijn et al. 2019), only a limited number of small-scale studies in broad-acre crops have demonstrated that these can be successfully introduced to complement or replace intensive farming practices (Horrocks et al. 2010; Macfadyen et al. 2014). Even fewer have shown that this is possible without concomitantly affecting crop profitability, and very few studies have been co-designed between practitioners and researchers to explicitly contribute to bridge the gap between science and practice (Caron et al. 2014; Cadotte et al. 2017; Hölting et al. 2022; Kurle 2024).

Here, we report empirical evidence of the ecological and economic benefits of sustainable agricultural practices from two co-designed field trials conducted across two years in collaboration with commercial grain producers in Australia. Our study is situated within the agricultural extension framework of on-farm demonstration sites – the use by researchers of methods and technologies that can be subsequently advised by agronomists and applied by farmers to generate positive practice change (Adamsone-Fiskovica and Grivins 2022). We tested whether a major pest – the redlegged earth mite – can be effectively managed by complementing conventional practices with targeted sustainable agriculture practices, including biological control and biofertiliser supplementation. Specifically, we used counterfactual treatments to disrupt the sustainable practices currently employed by each of the farmers to understand the effects of having had managed the system with conventional broad-spectrum insecticides and synthetic fertilisers. Additionally, we harnessed the power of precision agriculture to obtain fine-grained, paddock-level yield data, which we used to conduct cost-benefit analyses to assess whether the sustainable practices remained profitable. As such, our study contributes much needed evidence that sustainable agricultural practices can be applied to complement traditional approaches. We identified key success and limiting factors for the implementation of sustainable pest management practices in broad- acre grain systems, providing a framework for future research and extension, and highlighting the value of research/practice co-designed experiments and the merits of demonstration sites for agricultural extension.

## Materials and Methods

### Model pest: the redlegged earth mite

The redlegged earth mite (RLEM), *Halotydeus destructor* (Tucker), is one of the most economically important and destructive pests of winter pastures and grain crops in Australia (Ridsdill-Smith et al. 2008).

The RLEM occurs in the Mediterranean-type climate regions of southern Australia and is active in the cooler periods from April to November, typically undergoing three or four generations annually (Maino et al. 2024). The mites survive the hot summer months as diapause eggs in the cadaver of adult females on the soil surface, hatching out the following autumn after exposure to adequate rainfall and cooling temperatures (McDonald et al. 2015). Escalating issues with insecticide resistance have made control of this species using traditional chemical approaches challenging (Arthur et al. 2021).

### On-farm trial experimental design

The study was conducted in Victoria, Australia in two cropping paddocks (i.e. the demonstration sites), which were within farms located near Tennyson (36°18′ S, 144°26′ E) and Stoneleigh (37°36′ S, 143°13′ E), which were 160 and 72 ha, respectively. Within each we established three 10 ha experimental treatment plots, labelled Conventional, Conventional Plus and Novel (Tennyson) and Conventional, Novel and Novel Plus (Stoneleigh).

The Novel plots at both sites demonstrate the ensemble of sustainable practices that have been used by each farmer for multiple cropping seasons. At Tennyson, this comprises: (1) a plant nutrient regime focused on biological fertilisers but complemented with tactical supplements of synthetic fertilisers; and (2) pest and disease management based on biological control (i.e. seed and crops are not treated with agrochemicals) (Table 1). At Stoneleigh, it comprised a pest management regime based on biological control (Table 2). In both cases, the Novel plots are effectively a 10-ha subsample of the rest of the paddock.

**Table 1.**
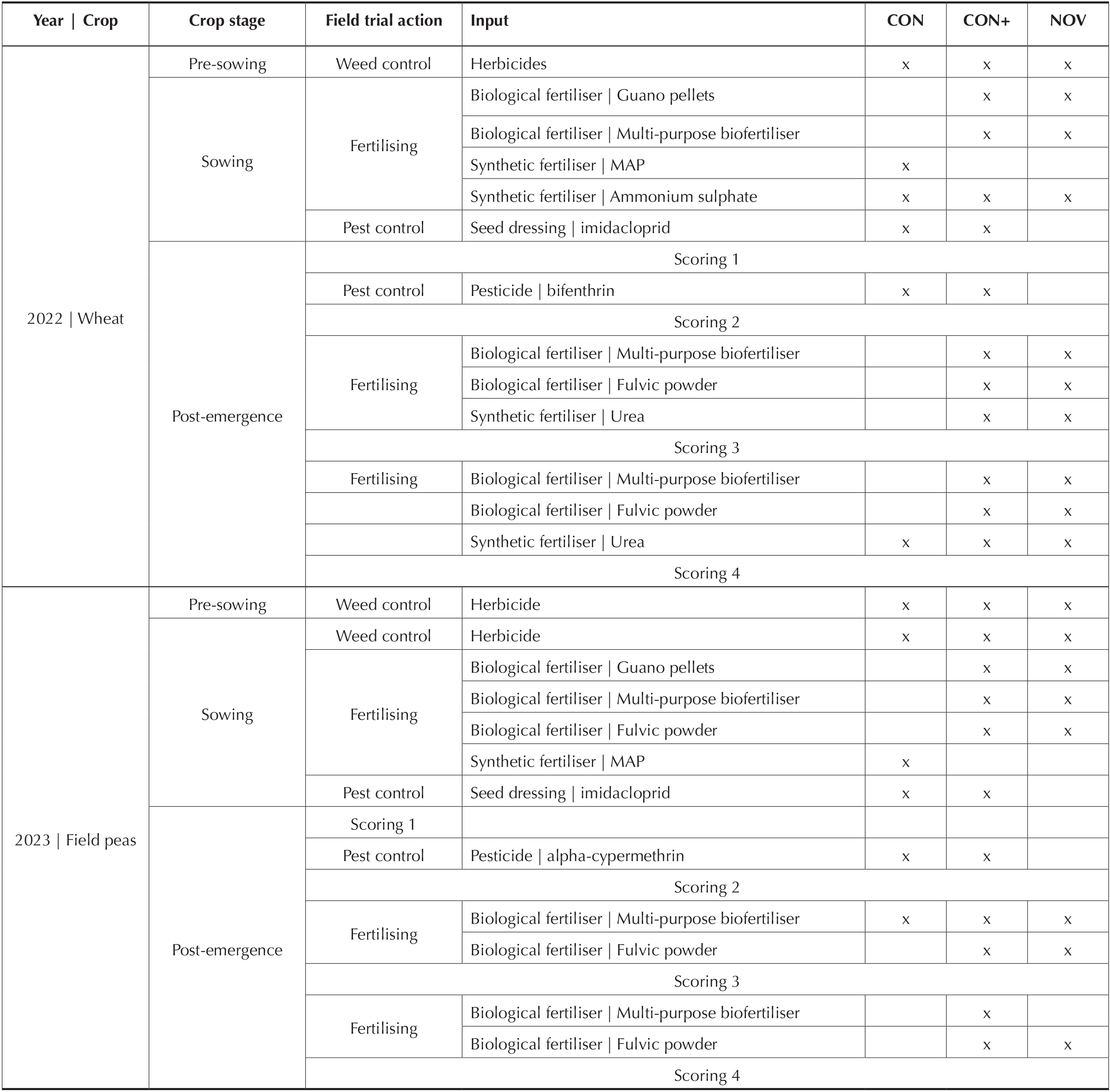
Inputs by treatment for the Tennyson site. The year, crop, crop stage and field trail action for each input are indicated for each input.

**Table 2.**
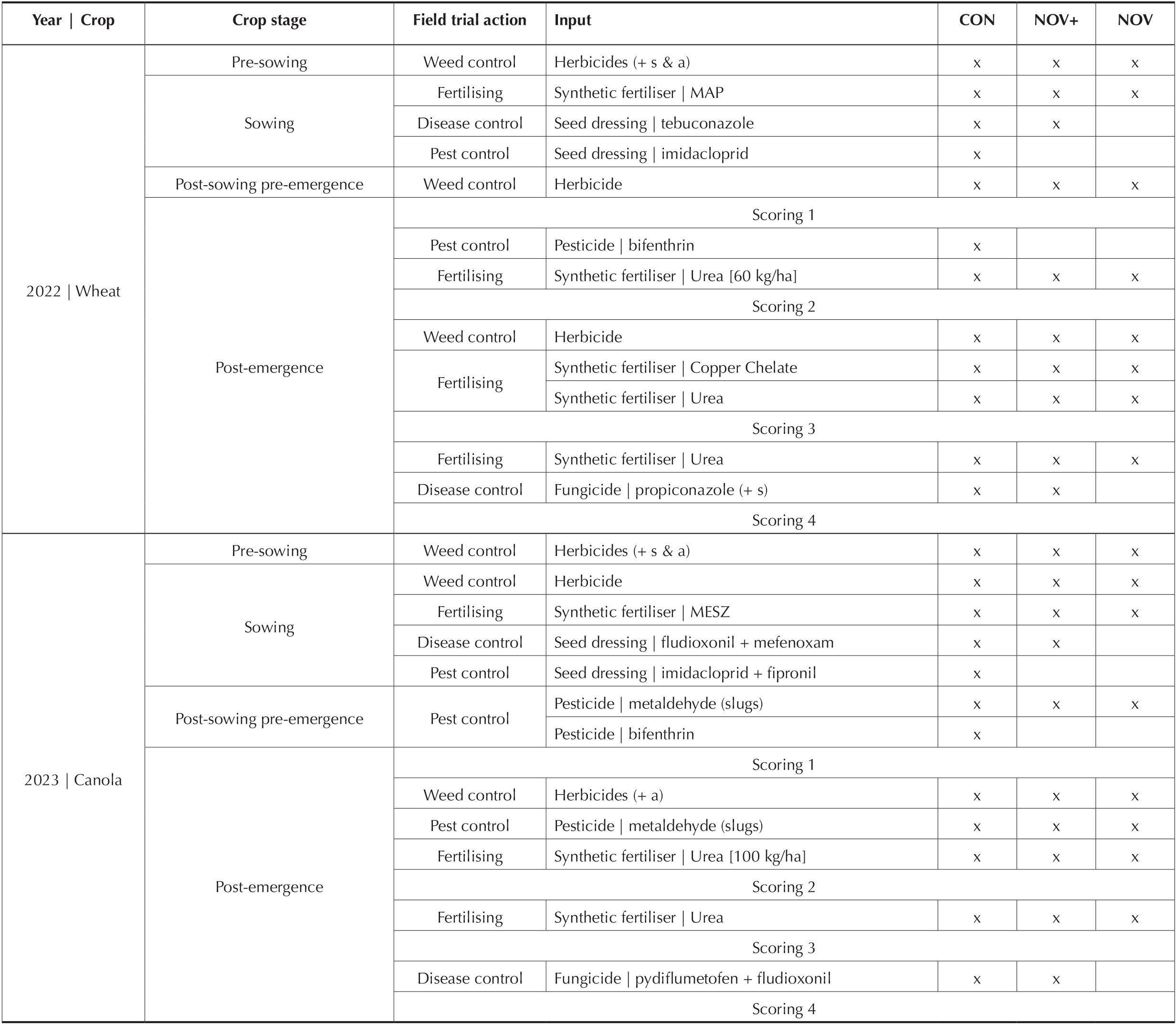
Inputs by treatment for the Stoneleigh site. The year, crop, crop stage and field trail action for each input are indicated for each input. a: adjuvant; s: surfactant.

The Conventional plots at both sites are intended to counterfactually demonstrate the effect of having had managed the paddocks without the sustainable practices currently employed at the Novel plots. This was achieved by: (1) a plant nutrient regime based on synthetic fertilisers (Tennyson; Table 1), and (2) a pest management regime based on chemical control (i.e. seed and crops treated with agrochemicals) (Tennyson and Stoneleigh; Tables 1 & 2).

The Plus plots are also intended as experimental counterfactuals. At Tennyson, the Conventional Plus plot demonstrates the effect of having had managed the paddock with the plant nutrient regime used in the Novel plot, but the pest management regime used in the Conventional plot (Table 1) – as such, it is intended to disentangle the potential confounding effects of the plant nutrient treatments from the pest management treatments. At Stoneleigh, the Novel Plus plot demonstrates the effect of having had managed the paddock with the pest management regime used in the Novel plot, but also removing the use of agrochemicals to control fungal diseases (Table 2) – as such, it is intended to provide initial evidence of the effect of a future disease and pest management regime exclusively focused on biological control.

At Stoneleigh, two applications of metaldehyde baits were used across all treatment plots in 2023. This decision was made in conjunction with the farmer and agronomy advisor due to the large number of pest slugs present across the paddock and the high risk of canola seedling loss (Nash et al. 2007).

### Ecological surveys

We assessed pest and beneficial species four times at each site in each year (Tables 1 & 2). All scorings were conducted during the post-emergence crop stage and timed to key management periods. Scoring 1 took place 20-39 days after sowing and timed before the application of the foliar insecticides. Scoring 2 took place 69-82 days after sowing and timed 1-3 weeks after the application of foliar insecticides. Scoring 3 took place 98-112 days after sowing and timed after mid-season application of fertilisers. Scoring 4 took place 137-175 days after sowing and timed after all treatments had been applied. The exact scoring dates are given in Tables S1-S4.

At each scoring, we surveyed 15 points, which were configured in a zig-zagging pattern across the plots and separated from each other by at least 50 m. At each point, we placed a 50 x 50 cm square frame and used a Stihl™ Blowervac BG55 to sample mites and other arthropods on plants and the soil surface, which were transferred from a fine mesh placed over the end of the vacuum tube into a tray. We then counted the number of RLEMs and beneficial species known to predate on RLEM, including the French anystis mite (*Anystis wallacei*), snout mites (Family Bdellidae) and spiders (herein referred to as beneficials). While not the focus of the study, we also counted the number of blue oat mites (*Penthaleus spp*.) and lucerne fleas (*Sminthurus viridis*), two pests found co-occurring with RLEM. Other arthropod species collected in the vacuum samples (e.g. collembola) were excluded from this study. Where field identifications were uncertain, specimens were brought back to the laboratory for identification under a dissecting microscope.

### Economic data

#### Yield density

We obtained paddock-level yield data (t/ha) directly from the harvesters’ precision agriculture software, which was provided either directly as a spreadsheet (Stoneleigh) or shapefile (Tennyson). These were relatively large datasets, varying from approximately 25,000 to 525,000 yield datapoints. To derive treatment-level data, we intersected the yield data with the plots’ spatial footprints using the R package sf (Pebesma 2018). We then z-standardised the datasets and removed extreme outliers (-3 < z < 3).

#### Input and application costs

For each treatment, we documented the costs per ha invested to produce the corresponding crop at the given site and year. These included both input (e.g. seeds, fertilisers, pesticides) and application (e.g. sowing, harvesting) costs (Tables S1-S4).

#### Statistical modelling

##### Modelling the effect of treatment and scoring on pests and beneficials

To assess the effect of treatment and scoring period on the densities (individuals/m^2^) of pests and beneficials, we used generalised linear models (Kéry 2010). The spatio-temporally replicated scoring points were the unit of analysis for drawing inferences on density. The model is organised in a single level for the point-specific densities (N_*i*_), which was specified as:

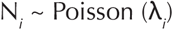

where λ_*i*_ is the density parameter of the Poisson distribution at scoring point *i*.

The linear predictor (log-probability scale) was specified as:

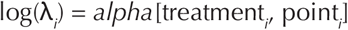

where treatment_*i*_ is an integer indexing the treatments; point_*i*_ an integer indexing the scoring point; and *alpha* the combined treatment/point fixed effects for the treatment/point combinations, which were specified as matrices and assigned to non-informative Normal priors with mean = 0 and precision = 0.001.

##### Modelling the effect of treatment on crop yield density

To assess the effect of trial treatment on crop yield (t/ha), we used general linear models (Kéry 2010). The spatially replicated yield points were the unit of analysis for drawing inferences on yield density. The model is organised in a single level for the point-specific densities (Y_*i*_), which was specified as:

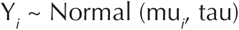

where mu_*i*_ and tau are the mean and precision parameters, respectively, of the Normal distribution, with tau = 1/sd^2^ and sd assigned to a Uniform (0, 100) prior.

The linear predictor was specified as:

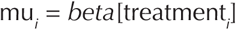

where treatment_*i*_ is an integer indexing the treatments; and *beta* the fixed effects for each treatment, which were assigned to non-informative Normal priors with mean = 0 and precision = 0.001.

##### Modelling the effect of treatment on gross profit margin

We used the yield by treatment estimates beta to calculate gross profit margins by treatments. The gross profit margin (GPM) equation was specified as:

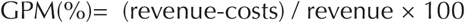

where costs (AUD/ha) are the costs by treatment for each site and year; and revenue (AUD/ha) calculated by multiplying the yield by treatment estimates *beta* (t/ha) by the selling price (AUD) of the corresponding crop at the given year. As these calculations were conducted within a Bayesian inference framework, the GPM estimates were derived with their full associated uncertainties.

##### Bayesian inference implementation

We used Markov Chain Monte Carlo simulations to draw samples from the parameters’ posterior distributions, with models implemented in JAGS (Plummer 2003), via the R package jagsUI (Kellner 2016). We used three chains of 5,000 iterations, discarding the first 500 iterations as burn-in. We assessed convergence by inspecting the chains and verifying that the values of the Gelman-Rubin statistic fell below 1.1 (Gelman and Hill 2007).

## Results

### Treatment effects on RLEM

At Tennyson, RLEM densities in the Conventional and Conventional Plus plots were low at the beginning of the wheat (2022) and field pea (2023) seasons (Scoring 1), remaining at very low densities throughout each season (Figs. 1 & 2; Table S5). In 2022, densities in the Novel plot were on average 6.3 times higher at Scoring 2 than 1, and 14.9 times higher at the end than beginning of the season (Figs. 1 & 2; Table S5). At Scoring 4, densities had increased to ∼320 mites/m2. In 2023, densities in the Novel plot were consistently higher than in the Conventional and Conventional Plus plots, but remained quite low (between ∼55 - 65 mites/m^2^). RLEM densities decreased sharply in the Novel plot at Scoring 4, which was likely due to mites going into summer diapause before this scoring event occurred (Maino et al. 2024) (Figs. 1 & 2; Table S5).

**Figure 1.**
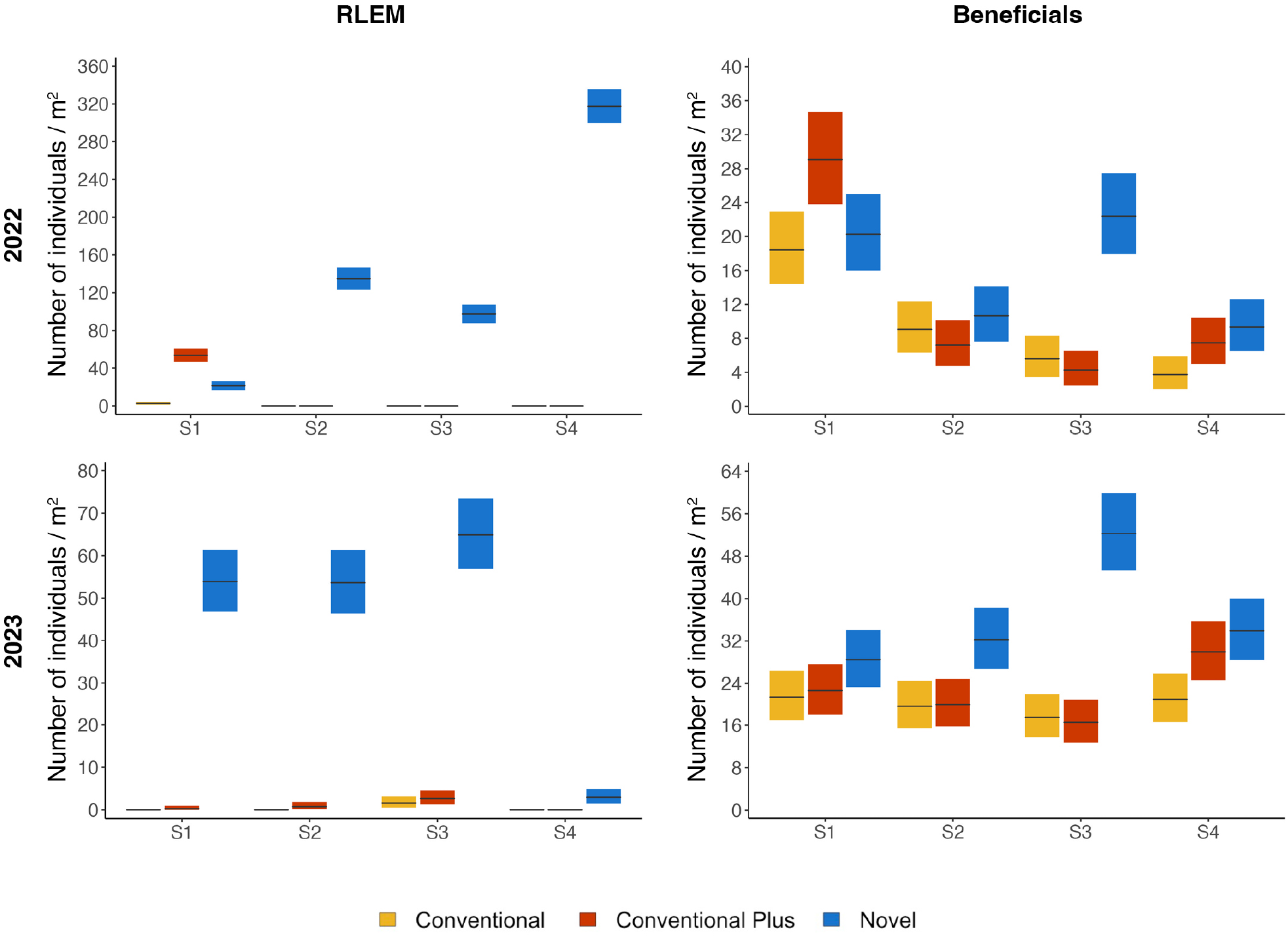
Redlegged earth mite and beneficial densities by year/crop (2022: wheat; 2023: field peas), experimental treatment (Conventional: yellow; Conventional Plus: red; Novel: blue) and scoring period (S1-S4) for the Tennyson demonstration site. The black horizontal lines represent the mean response and the coloured vertical bars the associated uncertainty (95% credible intervals).

At Stoneleigh, RLEM densities in the Conventional plot were low throughout the wheat (2022) and canola (2023) seasons (Figs. 3 & 4; Table S5). In the Novel plot, densities were also low throughout both seasons, increasing slightly (5.3 times on average) at Scoring 4 at the end of 2022 (Figs. 3 & 4; Table S5). Across 2022, densities were consistently higher in the Novel Plus plot; for example, densities were on average 4.1, 7.6 and 2.3 times higher at Scoring 2, 3 and 4, respectively, in the Novel Plus than Novel plot (Figs. 3 & 4; Table S5). In 2023, densities were consistently low in the Novel Plus plot, except at the end of the trial where densities had increased to ∼70 mites/m^2^ (Figs. 3 & 4; Table S5) – which was similar to the densities observed in the Novel Plus plot at the end of 2022.

In both sites and cropping seasons, and across all treatments, RLEM densities remained largely below the operational economic damage threshold of 1,000 mites/m^2^ in canola and 5,000 mites/m^2^ in wheat and field peas (Dunn and Miles 2000; Figs. 2 & 4).

**Figure 2.**
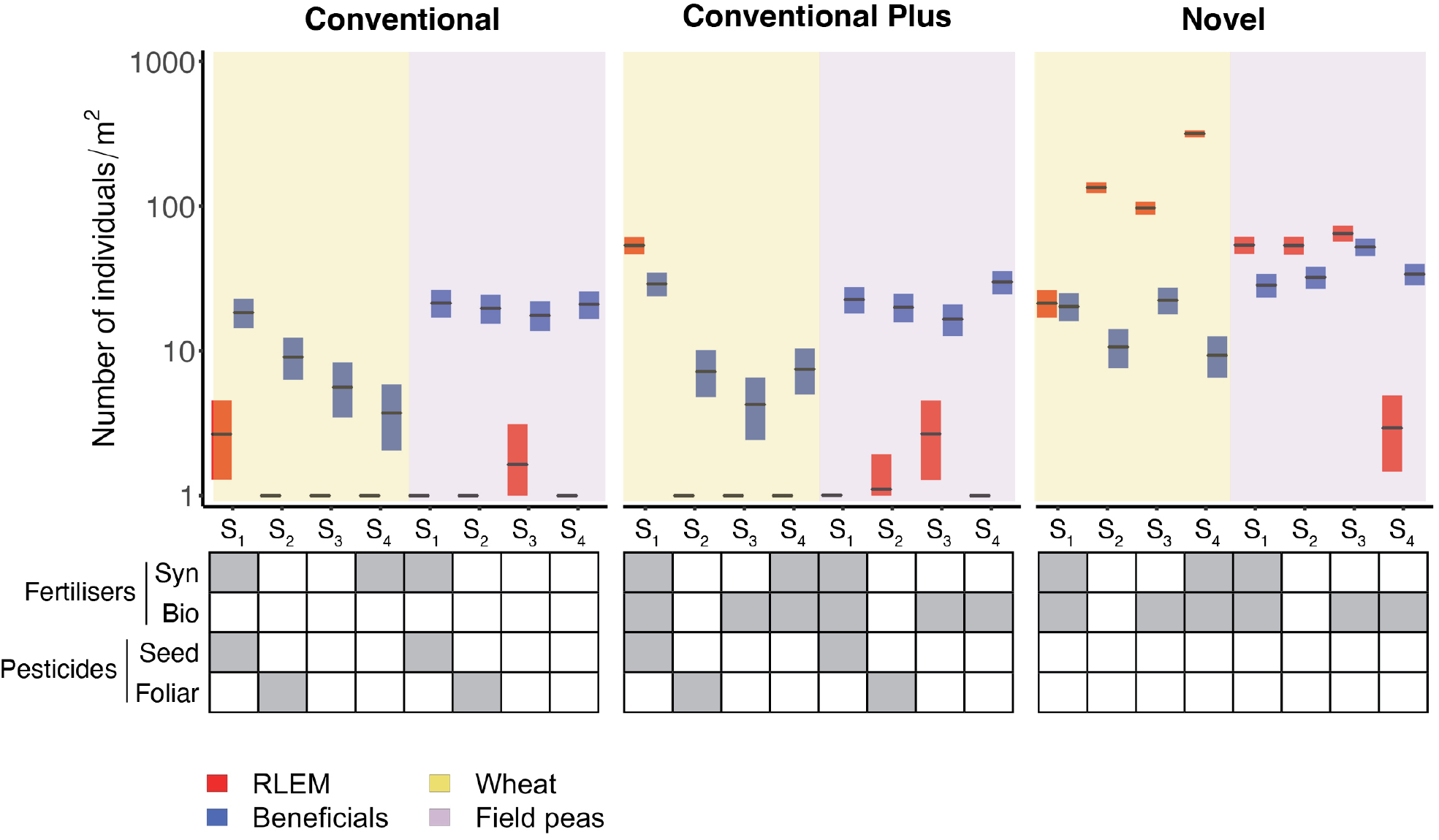
Side by side representation of redlegged earth mite (red bars) and beneficial (blue bars) densities by year/crop (wheat[2022]: yellow; field peas[2023]: light purple), experimental treatment and scoring period (S1-S4) for the Tennyson demonstration site. The matrix below each treatment summarises the fertiliser and insecticide inputs associated with each treatment and scoring. The black horizontal lines represent the mean response and the coloured vertical bars the associated uncertainty (95% credible intervals). Syn: synthetic; Bio: biological.

### Treatment effects on beneficials

At Tennyson, beneficial densities in 2022 decreased in the Conventional and Conventional Plus plots from Scoring 2 onwards (Figs. 1 & 2; Table S6). By Scoring 3, there was a marked difference between densities; the Novel plot contained ∼22 mites/m^2^, which was on average 4.0 and 5.2 times higher than the Conventional and Conventional Plus plots, respectively. A treatment effect was also observed at Scoring 4, where densities were on average 2.5 times higher in the Novel than the Conventional plot (Figs. 1 & 2; Table S6). In 2023, the Conventional and Conventional Plus plots showed consistent densities across the season, except for a slight increase at Scoring 4 as compared to Scoring 3 in the Conventional Plus plot (1.8 times on average; Figs. 1 & 2; Table S6). Beneficial densities were consistently higher in the Novel than the Conventional and Conventional Plus plots in 2023. As in 2022, this difference was greatest at Scoring 3, where the Novel plot contained ∼52 mites/m^2^, which was on average 3.0 and 3.2 times higher than the Conventional and Conventional Plus plots, respectively (Figs. 1 & 2; Table S6).

At Stoneleigh, beneficials densities in 2022 were consistently low in the Conventional plot across the season (Figs. 3 & 4; Table S6). Beneficials densities were also low in the Novel and Novel Plus plots at Scoring 1 and Scoring 2 but increased thereafter. By the end of the season, the Novel plot contained ∼12 mites/m^2^ and the Novel Plus plot contained ∼6 mites/m^2^, which was on average 11.5 and 6.0 times higher than the Conventional plot, respectively (Figs. 3 & 4; Table S6). In 2023, densities in the Conventional Plot were low at the beginning of the year and fluctuated throughout the season (Figs. 3 & 4; Table S6). Beneficial densities were also low in the Novel and Novel Plus plots at the beginning of 2023 but increased consistently through the season. By the end of the season, densities were on average 9.2 and 8.2 times higher in the Novel and Novel Plus plots, respectively, than at the beginning (Figs. 3 & 4; Table S6).

**Figure 3.**
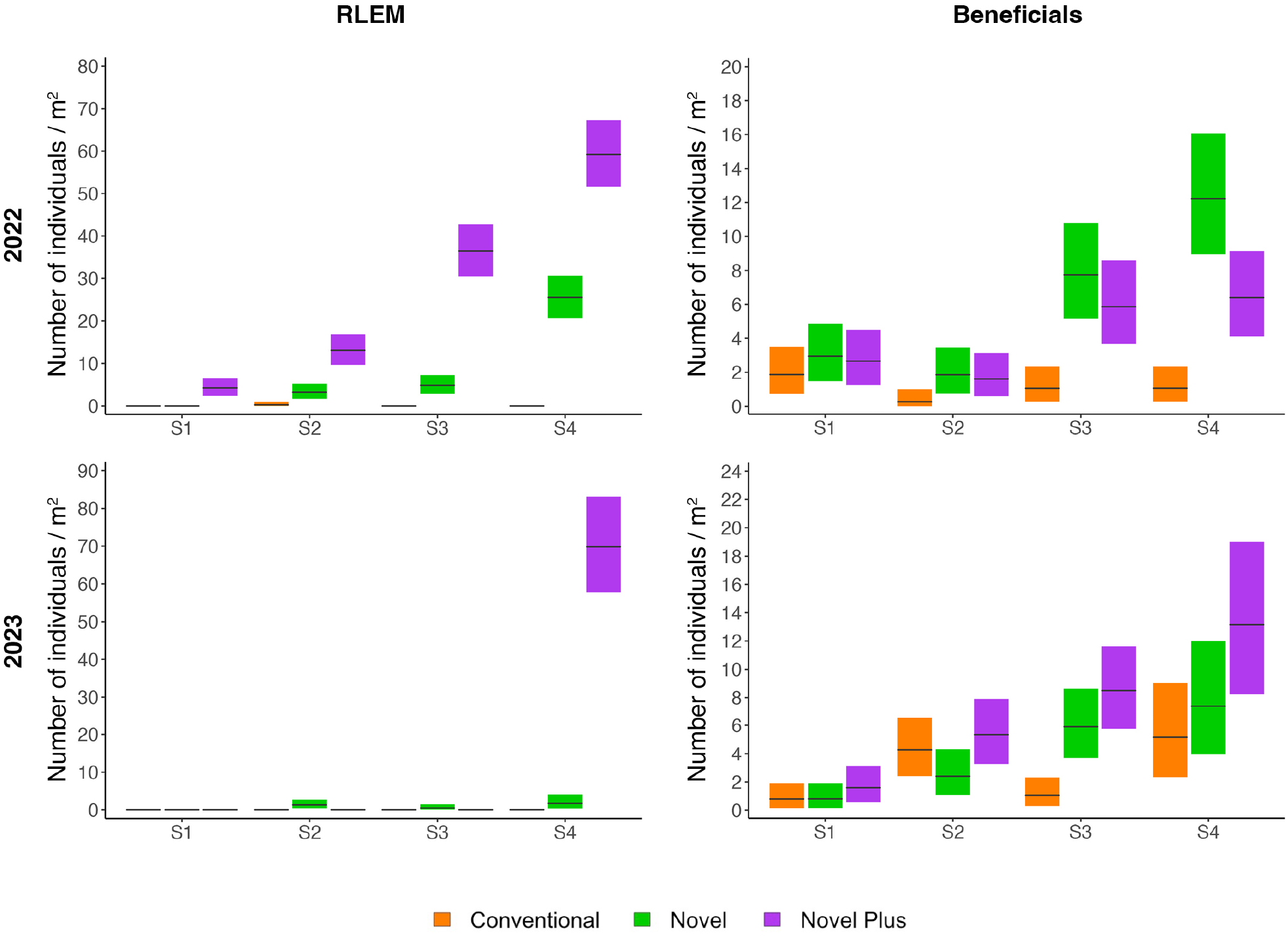
Redlegged earth mite and beneficial densities by year/crop (2022: wheat; 2023: canola), experimental treatment (Conventional: orange; Novel: green; Novel Plus: purple) and scoring period (S1-S4) for the Stoneleigh demonstration site. The black horizontal lines represent the mean response and the coloured vertical bars the associated uncertainty (95% credible intervals).

**Figure 4.**
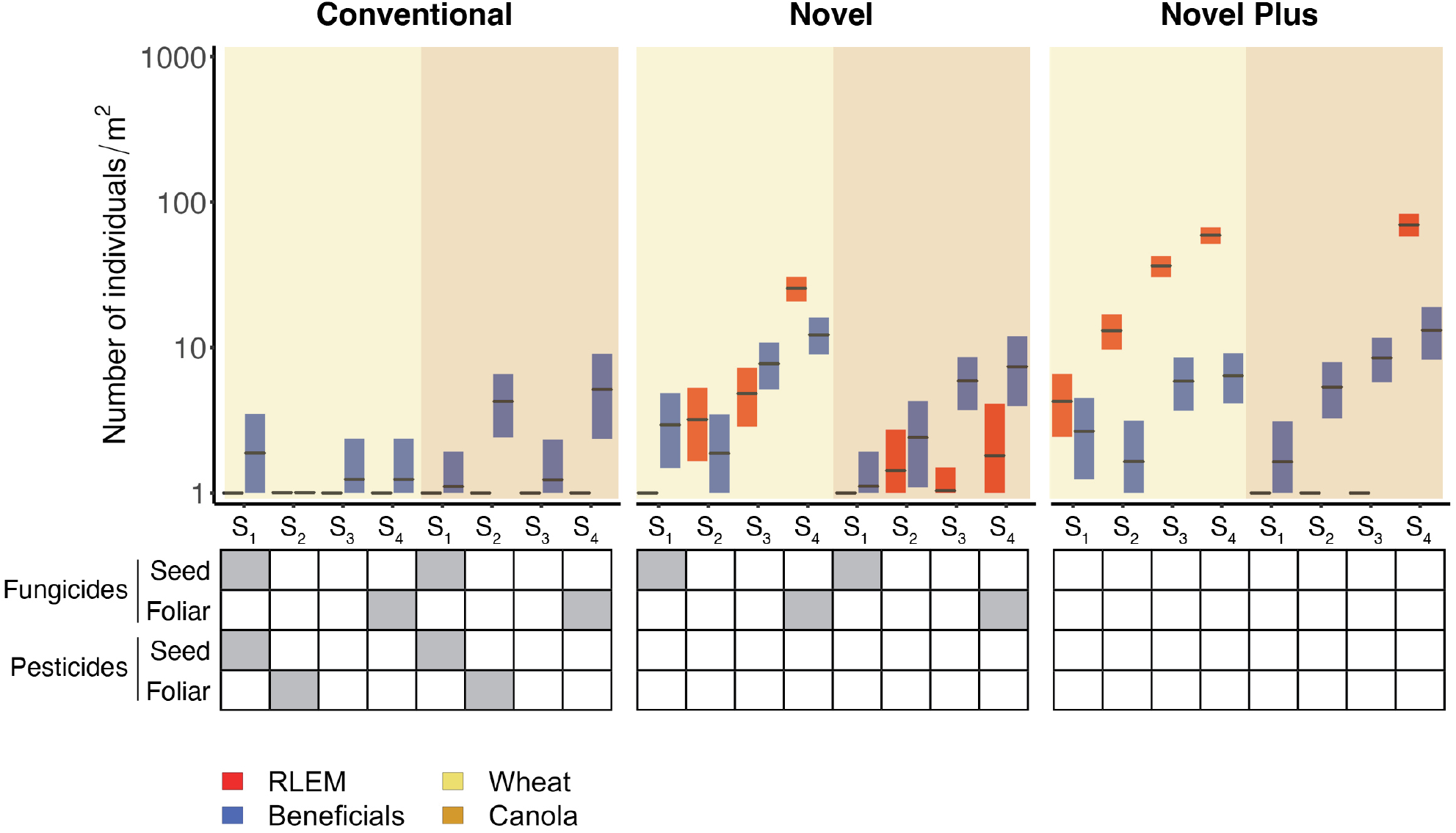
Side by side representation of redlegged earth mite (red bars) and beneficial (blue bars) densities by year/crop (wheat[2022]: yellow; canola[2023]: orange), experimental treatment and scoring period (S1-S4) for the Stoneleigh demonstration site. The matrix below each treatment summarises the fungicide and insecticide inputs associated with each treatment and scoring. The black horizontal lines represent the mean response and the coloured vertical bars the associated uncertainty (95% credible intervals).

### Treatment effects on other crop pests

At Tennyson in 2022, blue oat mite densities remained low (< 3.7 mites/m^2^) throughout the season across all treatments, only increasing at the end of the season in the Novel plot (densities were on average 7.7 times higher at Scoring 4 than Scoring 1; Fig. S1; Table S7). Lucerne flea densities were also relatively low at the beginning of 2022, increasing in the Conventional and Conventional Plus plots after the application of the foliar insecticide (Fig. S1; Table S8). Conversely, lucerne flea densities decreased in the Novel plot throughout the season; at Scoring 4, densities were on average 12.1 and 17.4 times lower than in the Conventional plot and Conventional Plus plot, respectively. In 2023, blue oat mite densities again remained very low, never exceeding an average density of >7 mites/m^2^, irrespective of treatment (Fig. S1; Table S7). Lucerne flea densities, however, were much higher in both the Conventional and Conventional Plus plots compared with the Novel plot at the beginning of 2023 (on average 3.1 and 3.8 times higher in Conventional and Conventional Plus plots, respectively). Lucerne flea densities decreased in all treatments after Scoring 1 and remained low thereafter (Fig. S1; Table S8).

At Stoneleigh, blue oat mite densities were very low at the beginning of 2022 and 2023 across all treatments (Fig. S2; Table S7). In 2022, blue oat mite densities either remained very low in Conventional plot or increased in the Novel (10.1 times on average) and Novel Plus (8.4 times on average) plots between Scoring 2 and Scoring 4 (Fig. S2; Table S7). In 2023, blue oat mite densities fluctuated throughout the season. By Scoring 4, the Conventional plot had considerably more blue oat mites than the Novel and Novel Plus plots (Fig. S2; Table S7).

### Effects of treatment on yield and gross profit margins

At Tennyson, wheat yield between the Conventional and Novel plots in 2022 were not statistically different from each other, with both treatments yielding on average 1.77 and 1.83 times lower, respectively, than the Conventional Plus plot (Fig. 5; Table S9). Similarly, gross profit margins between the Conventional and Novel plots were not statistically different from each other, with both treatments resulting in profit margins that were on average 3.56 and 3.17 times lower, respectively, than those estimated for the Conventional Plus plot (Fig. 5; Table S9). In 2023, field peas yield was higher in the Conventional plot than in the Novel plot, which yielded slightly higher than the Conventional Plus plot. Yield in the Novel plot was on average 1.09 times lower than the Conventional plot and 1.09 times higher than the Conventional Plus plot (Fig. 5; Table S9). The resulting gross profit margin was 1.14 times higher in the Conventional than in the Novel plot, while the Conventional Plus plot resulted in gross profit margins that were 1.79 and 2.05 times lower than those estimated in the Novel and Conventional plots, respectively (Fig. 5; Table S9).

**Figure 5.**
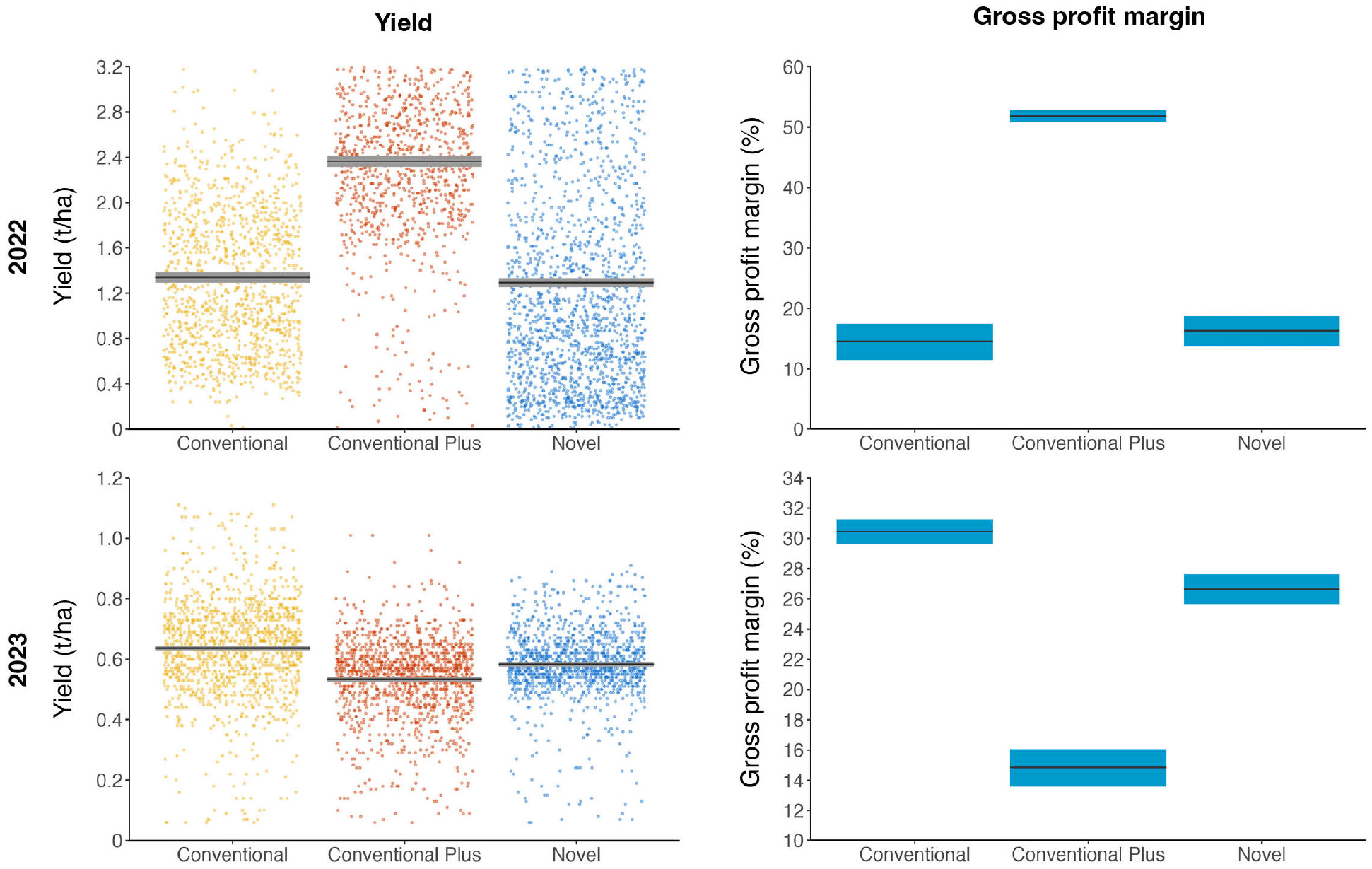
Yield and gross profit margins by year/crop (2022: wheat; 2023: field peas) and experimental treatment for the Tennyson demonstration site. The black horizontal lines represent the mean response and the grey (yield) and blue (gross profit margin) vertical bars the associated uncertainty (95% credible intervals). The coloured dots in the yield plot represent the spread of yield data obtained from the combine harvesters’ precision agriculture software (Conventional: yellow; Conventional Plus: red; Novel: blue).

At Stoneleigh in 2022, wheat yield was 1.03 times higher in the Conventional than in the Novel plot and 1.13 times higher in the Conventional than in the Novel Plus plot (Fig. 6; Table S9). Gross profit margin on the other hand was 1.03 times higher in the Novel Plus than in the Conventional plot and 1.08 times higher in the Novel Plus than in the Novel plot (Fig. 6; Table S9). In 2023, canola yield was 1.05 times higher in the Conventional than in the Novel plot and 1.09 times higher in the Conventional than the Novel Plus plot (Fig. 6; Table S9). The gross profit margins, however, were 1.03 times higher in the Novel Plus than the Conventional plot and 1.05 times higher in the Novel Plus than in the Novel plot (Fig. 6; Table S9).

**Figure 6.**
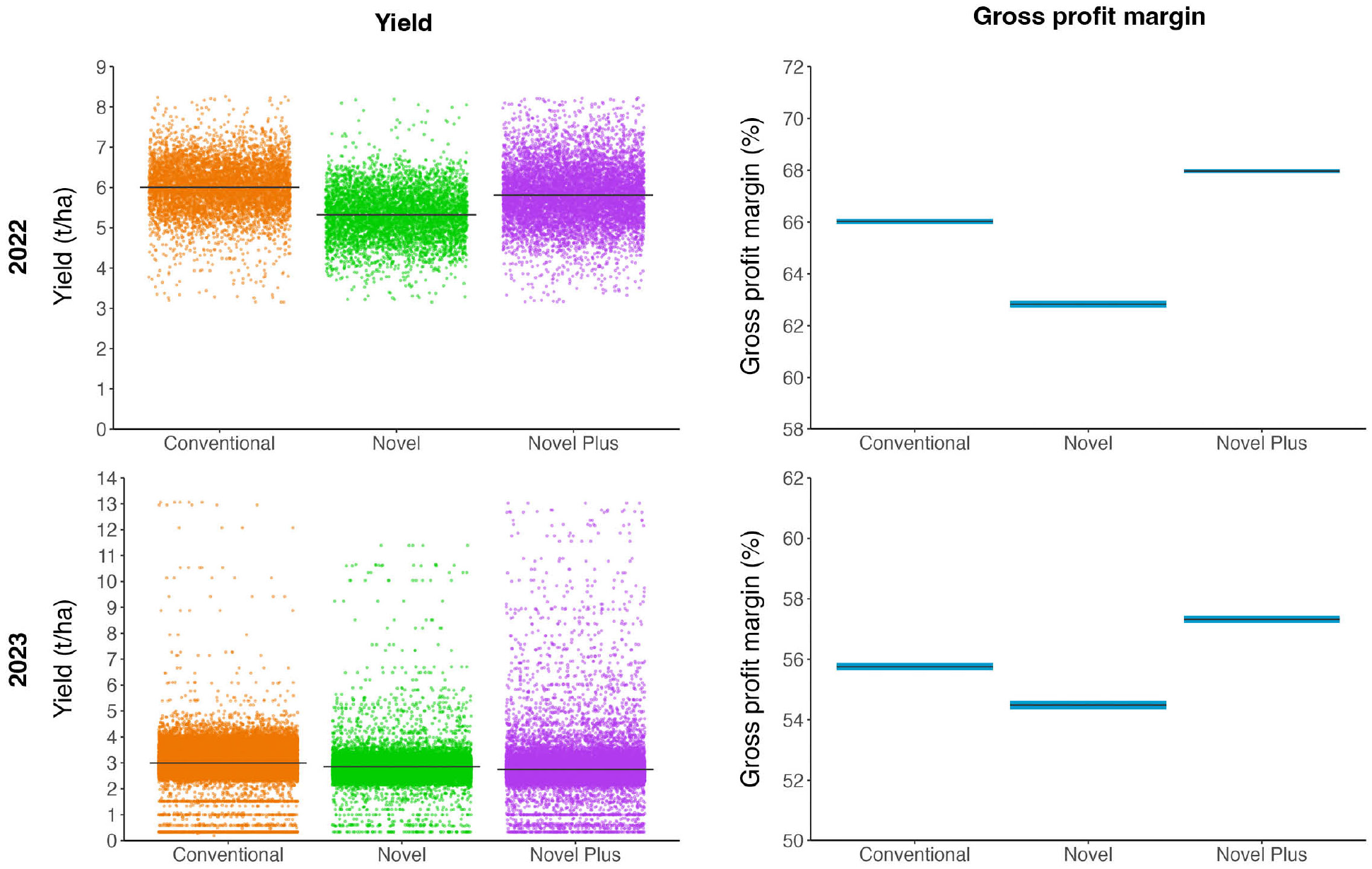
Yield and gross profit margins by year/crop (2022: wheat; 2023: canola) and experimental treatment for the Stoneleigh demonstration site. The black horizontal lines represent the mean response and the grey (yield) and blue (gross profit margin) vertical bars the associated uncertainty (95% credible intervals). The coloured dots in the yield plot represent the spread of yield data obtained from the combine harvesters’ precision agriculture software (Conventional: orange; Novel: green; Novel Plus: purple).

## Discussion

In this study, we provide new empirical evidence of the ecological and economic benefits of sustainable agricultural practices. We show that populations of the RLEM – one of the most economically important pests of winter pastures and grain crops in Australia – remained below economic thresholds across a range of crops in Conventional plots managed with broad-spectrum insecticides and synthetic fertilisers and in Novel plots managed with biological control and biofertiliser supplementation, indicating effective pest control in both approaches. Our findings therefore contribute much needed evidence that sustainable agricultural practices can achieve comparable pest control outcomes to chemical-centric practices, addressing a key concern for farmers considering adoption.

Additionally, the Novel and Novel Plus treatments consistently supported higher densities of beneficial species, indicating greater potential for natural pest control. This aligns well with previous studies reporting on the adverse effects of broad-spectrum insecticides on beneficial species – including the neonicotinoid seed treatments and synthetic pyrethroid foliar sprays that were tested here (Desneux et al. 2007; Schulz et al. 2021; Mata et al. 2024). Beyond impacting the natural enemies directly responsible for controlling pest populations, broad-spectrum insecticides can also harm other non-target beneficial species, triggering negative feedback loops that further affect yields and profits. For example, the application of synthetic pyrethroids to control RLEM and other crop pests can result in the secondary flare up of other pests, such as the lucerne flea (Michael 1991; Roberts et al. 2009). Indeed, at the Tennyson site in 2022, where lucerne fleas were present, densities increased sharply in the Conventional and Conventional Plus plots after a single application of the synthetic pyrethroid bifenthrin. Additionally, pesticide residues in the pollen and nectar of canola and other oilseed rape cultivars can compromise the fitness and survival of managed and wild bees (Rundlof et al. 2015; Tosi et al. 2022), affecting their capacity to boost yield quantity and quality (Catarino et al. 2019; Perrot et al. 2024). Moreover, emerging research is shedding light on the harmful effects of pesticides on the structural and functional diversity of soil, plant, and arthropod-associated microbiomes, with implications for crop performance (Edlinger et al. 2022; Compant et al. 2024). We hope our study serves as a further incentive for future research into harnessing the ecological benefits provided by beneficial organisms in agriculture.

Importantly, crop yields and gross profit margins were similar between the Conventional and Novel treatments across all trials, although at Stoneleigh in 2022, the wheat yield and resulting gross profit margin were noticeably greater when employing a pest management regime that relied on pesticide inputs. The average yield was 6.01 t/ha and the gross profit margin 66.03% in the Conventional plot compared with 5.32 t/ha and 62.83% in the Novel plot. Nonetheless, our findings are encouraging and support previous studies demonstrating that sustainable agriculture practices aimed at reducing agrochemical use can be economically competitive (Macfadyen et al. 2014; Lechenet et al. 2017). The landmark study by Lechenet and colleagues (2017) epitomises this idea – using data from over 900 non-organic commercial farms they found that in as much as 77% cases it was possible to reduce pesticide use while maintaining high productivity and profitability. Some of our yield and gross profit margin results, however, are more challenging to explain and require a more nuanced interpretation, particularly when considering the Novel Plus and Conventional Plus treatments. For example, at Tennyson in 2022, the Conventional Plus plot had higher wheat yield and gross profit margin than the Conventional and Novel plots. This suggests the combination of a biofertiliser-based plant nutrient regime and a pesticide-based pest management regime can be highly profitable, although this was clearly not the case in 2023 when growing field peas, where the Conventional Plus plot had the lowest yield and gross profit margin. Additionally, at Stoneleigh in 2022, the Novel Plus plot had considerably higher wheat yield and gross profit margin than the Novel plot – despite these two treatments being the same except for the fungicide applications in the Novel plot. The reason(s) for greater yield in the absence of fungicides remains unclear but may suggest non-target impacts of fungicides on arthropods and/or soil biota with detrimental consequences for nutrient recycling, mycorrhizal symbioses and biological pest control (Hage-Ahmed et al. 2018; Fernández et al. 2015). These results point to the complexity and high dimensionality of agroecosystems and highlights the benefit of conducting studies such as this, which enable intricate interactions between alternative farming practices to be explored – as opposed to short-term small plot trials, which typically prohibit the study of invertebrate ecology.

A particular strength of this study was the research/ practice co-design approach, which ensured the research questions and methods were closely aligned with the priorities and realities of the participating farmers. This methodology strengthens the practical applicability of the results while encouraging wider adoption, aligning well with (1) calls to untap the power of co-design to bridge the gap between science and practice (Caron et al. 2014; Cadotte et al. 2017; Hölting et al. 2022; Kurle 2024); and (2) mandates from funding bodies requiring researchers to work closely with practitioners to improve the likelihood that findings are properly translated into actionable management recommendations and, in time, potentially also into strategic plans and long-term policies. Indeed, beyond the extensive on-ground knowledge and economic in-kind contributions provided by the participating farmers and their agronomy advisors, the insights gained through this study have contributed to industry-relevant management recommendations for the control of RLEM in Australian grain crops.

For one of the participating farmers, the study reaffirmed their decision to invest in the health of the crop through biological additives rather than relying on conventional insecticides to control arthropod pests. For them, this highlights the opportunity to shift seed treatment strategies from neonicotinoid to biological and mineral nutrition seed coatings, which can promote seed vigour during germination and improve the rhizome’s ability to form strong symbiotic relationships with the soil microbiome (Mor et al. 2019; Yu et al. 2024). For them this exemplifies their current management approach focused on the relationships between plant health and pest damage, whereby plant sap analysis is used throughout the season to measure nutrient levels, allowing for targeted fertiliser applications to address potential deficiencies. Importantly, they pointed out that when making decisions on whether to prioritise nutrition over prophylactic chemical control during critical crop stages careful attention needs to be paid to environmental factors (e.g. frosts) that may adversely affect the capacity of the crop to deter pests without chemical assistance, highlighting situations where pesticides might be warranted to keep RLEM under economic thresholds. Indeed, both participating farmers have a long history of implementing sustainable practices with minimal to no insecticide use, including in the paddocks used in this study. It was both surprising and encouraging to find that, at the beginning of the trials, RLEM densities were very low despite the absence of insecticide applications in these paddocks for several years. Furthermore, RLEM densities remained consistently low and well below economic threshold levels (Dunn and Miles 2000) throughout the two-year study period, supporting the efficacy of the farming approaches being employed on each farm.

The participating farmers and advisors also emphasised how their knowledge about RLEM in-field identification, monitoring, lifecycle and natural enemies reinforced their ability to manage this pest proficiently. One of the advisors, for example, called attention to how they stress to their clients how sowing canola in paddocks that have just come out of pasture would not work unless strong pesticide applications against RLEM are used, as monitoring experience indicates that the pest is known to build up dense populations in pastures, a system that is, incidentally, well known for hosting RLEM insecticide-resistant populations. This latter example accentuates the key role that RLEM extension efforts play in supporting Australian farmers and advisors exposed to this difficult pest.

We recognise that our study allows for several areas of improvement, opening the door to new opportunities for future research. First, the investigated response variables were modelled with appropriately replicated within-plot data, yielding robust inferences that allowed for sound statistical interpretations. The replication at the paddock/farm level, however, was limited to the two demonstration site case studies reported here, which allowed us to investigate only three crops grown in two different subregions of temperate Australia. While caution must be taken in generalising our results to other crops and regions under different climates, the knowledge acquired from these specific farms can inform the selection of treatments and spatial scales for subsequent research across a wider range of conditions. Importantly, the study’s data and statistical inferences may be combined with that from other similar studies – for example through meta-analysis– to increase the accuracy of the results and strengthen confidence that the observed outcomes are caused by the researched practices (Ockendon et al. 2021). Another important limitation was that, while we provide observational evidence of the effects of reducing pesticide-use on supporting higher densities of beneficial species, additional experimental techniques would have been needed to fully evidentiate the capacity of beneficials to control pests. As Macfadyen and colleagues (2015) point out, assessing the impact of beneficials on pests does not need to be either difficult, time consuming or cost prohibitive. As them, we urge future field studies to embed in their design measurements capable of quantifying the magnitude by which beneficials are suppressing the targeted pest. Despite these limitations, our study can be positioned as a first step towards designing more comprehensive and replicated experiments in the future.

This study represents one of the few examples of co-designed research conducted in real-world farming systems that have sought to meet the challenge of providing empirical evidence of the ecological and economic benefits of sustainable agricultural practices. Additionally, we did not observe considerable increases in the abundance of RLEM when we counterfactually switched to a pest management strategy based on insecticide inputs over two consecutive years, pointing to the resilience of the agroecosystems each farmer had ‘built up’ over many years of implementing sustainable practices. The evidence that both farmers were on the right trajectory with their ‘novel’ practices enriches our awareness of how sustainable agricultural practices can be achieved on a large scale while remaining commercially viable. Our findings also underscore the opportunity to effectively manage redlegged earth mites with reduced reliance on broad-spectrum insecticides, thereby supporting resistance management practices that have long been recommended for this pest. We hope our study encourages broader adoption of sustainable agriculture practices and sparks further collaboration between farmers, advisors and scientists.

## Acknowledgements

This research was funded by the Grains Research and Development Corporation, grant number CES2010- 001RTX. We gratefully acknowledge the significant contribution of Adriana Arturi and the support provided by Lachlan McDougall. We thank Ary Hoffmann, Gary McDonald, Chris Dunn and Matt Page for their technical contributions, and Greg Toomey and Tom De Mattia for their generous assistance in compiling the input and application costs used to estimate gross profit margins. We acknowledge the Dja Dja Wurrung, Taungurung, Yorta Yorta, Eastern Maar, Wadawurrung, Woi wurrung and Boon wurrung / Bunurong peoples as the Traditional Custodians of the land and waterways on which this research took place – we pay our respects to their Elders, past, present and emerging.

## Author Contributions

James Maino and Paul Umina conceptualised and acquired funds for the project; Luis Mata, James Maino, Grant Sims, Craig Drum and Paul Umina designed the methodology; Luis Mata, Leo McGrane, James Maino, Grant Sims, Craig Drum and Paul Umina conducted the research; Luis Mata curated and analysed the data; Luis Mata and James Maino administered the project; Luis Mata, James Maino and Paul Umina supervised staff; Grant Sims and Craig Drum provided experimental resources and equipment; Luis Mata and James Maino conceptualised and developed visualisations; Grant Sims and Craig Drum provided management recommendations; Luis Mata led the writing of the manuscript. All authors contributed critically to the drafts and gave final approval for publication.

## Data availability statement

Field data and codes to reproduce models and plots are already published and publicly available in Zenodo: https://doi.org/10.5281/zenodo.13958616

---

**Figure S1.**
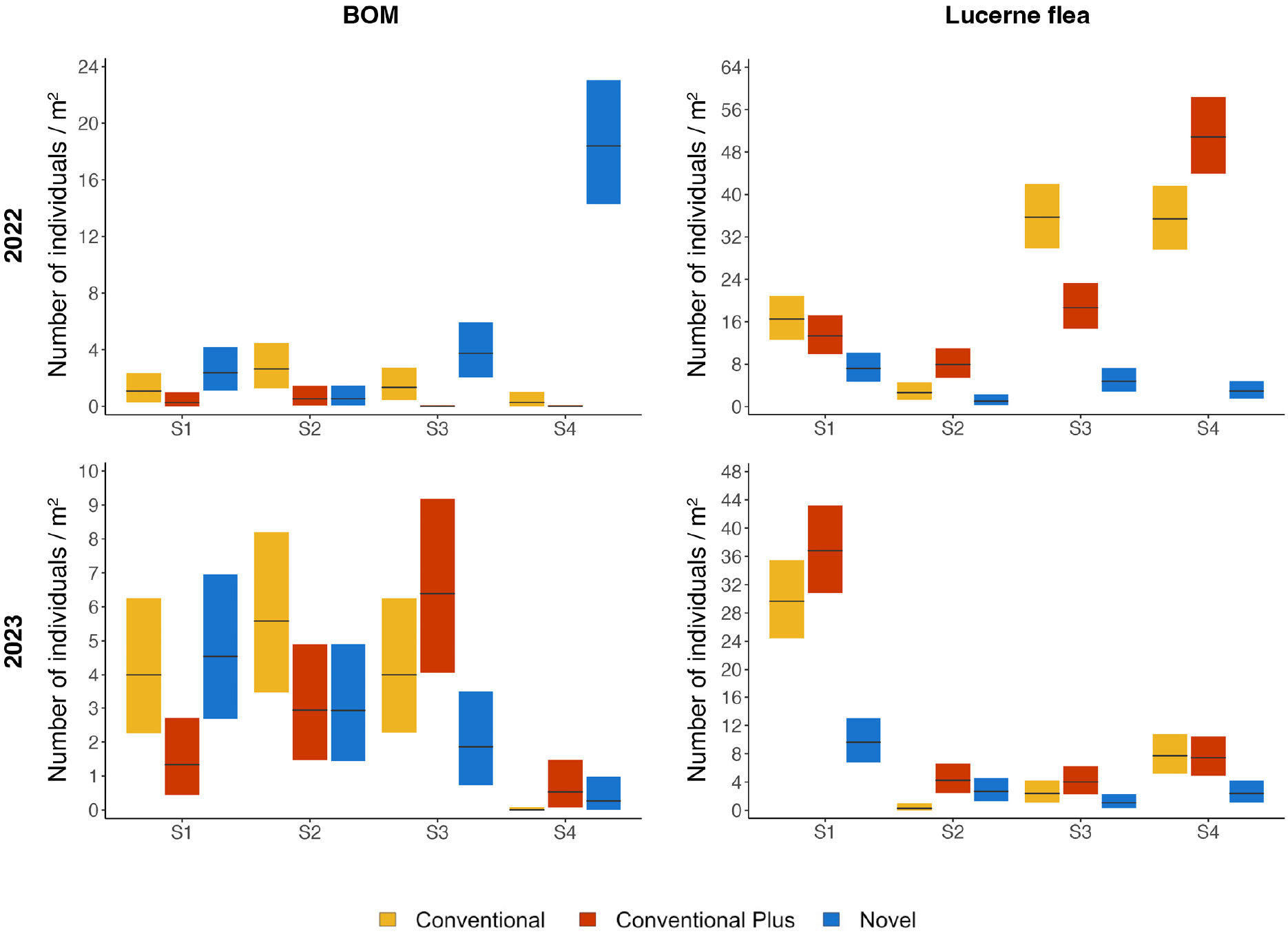
Blue oat mite and lucerne flea densities by year/crop (2022: wheat; 2023: field peas), experimental treatment (Conventional: yellow; Conventional Plus: red; Novel: blue) and scoring period (S1-S4) for the Tennyson demonstration site. The black horizontal lines represent the mean response and the coloured vertical bars the associated uncertainty (95% credible intervals).

**Figure S2.**
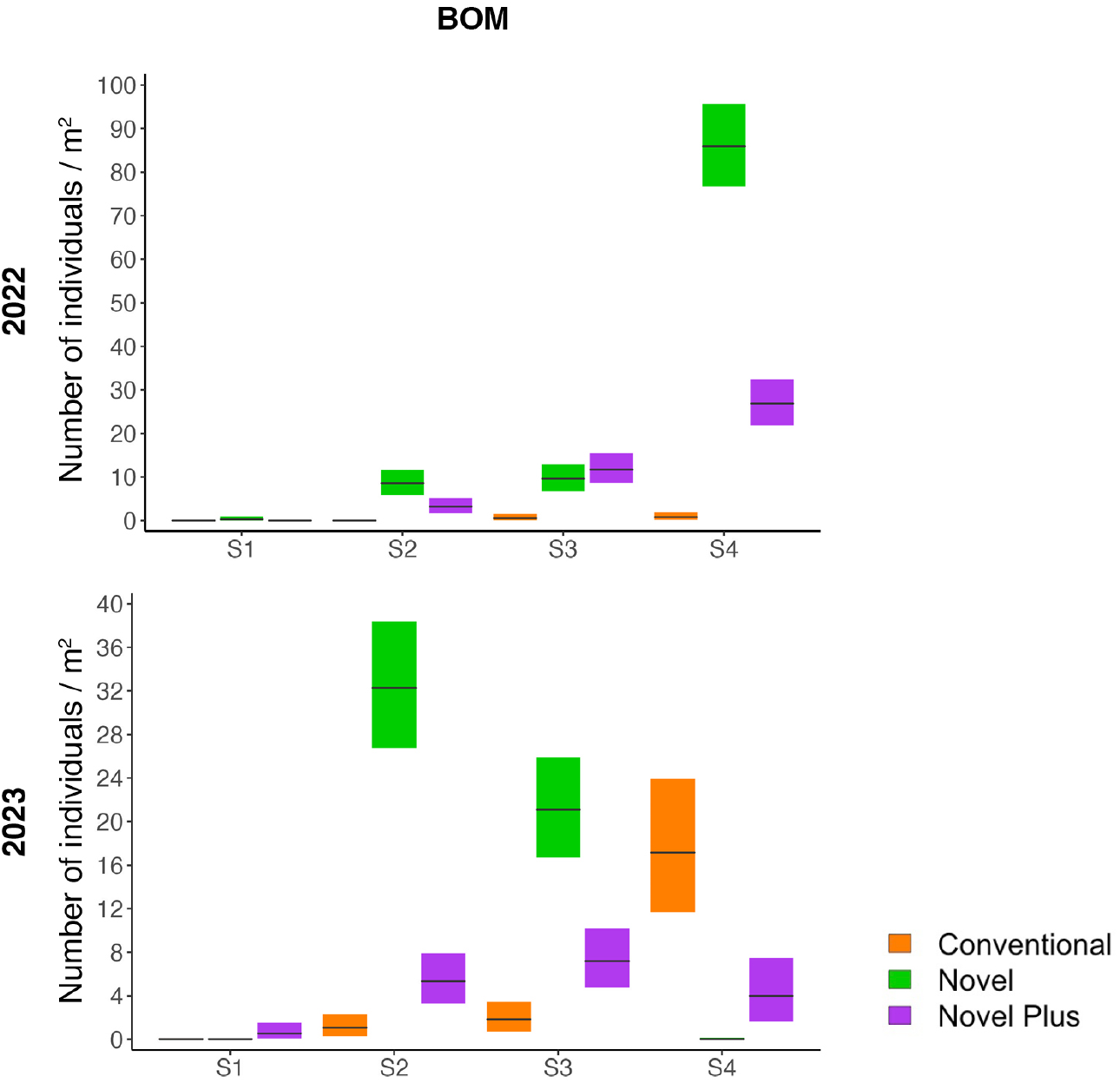
Blue oat mite densities by year/crop (2022: wheat; 2023: canola), experimental treatment (Conventional: orange; Novel: green; Novel Plus: purple) and scoring period (S1-S4) for the Stoneleigh demonstration site. The black horizontal lines represent the mean response and the coloured vertical bars the associated uncertainty (95% credible intervals).

